# Dietary and behavioral inferences from dental pathology and non-masticatory wear on dentitions from a British medieval town

**DOI:** 10.1101/222091

**Authors:** Ian Towle, Carole Davenport, Joel D. Irish, Isabelle De Groote

**Author notes:** **Contact details**: Ian Towle, Room 352, James Parsons Building, Byrom Street, Liverpool, L3 3AF, 07880330161.

## Abstract

Dental pathology and wear data can provide valuable insights into diet, cultural practices, and the health of populations. In this study, various dental pathologies and types of wear were recorded for 41 individuals (914 permanent teeth), excavated from the medieval cemetery of St. Owens Church in Southgate Street, Gloucester. Teeth were studied macroscopically with a 10x hand lens to confirm the presence of specific pathologies. Relatively high rates of antemortem chipping on the anterior teeth, and the presence of maxillary central incisor notches, suggested that the Gloucester population commonly used their teeth for non-masticatory activities. Abscessing and antemortem tooth loss fell within previously reported ranges for British medieval sites (2.6% and 6% respectively). However, the sample exhibits extremely high levels of carious lesions and calculus. Nearly 24% of teeth have at least one carious lesion, and the presence of calculus was recorded in 74% of teeth within the sample. Overall caries frequency is similar to sites from later time periods. This frequency may reflect Gloucester’s location as a large port town. Remains from the same area, but the earlier Roman period, also shows high rates of both caries and calculus, suggesting a continuation of consuming certain cariogenic foods is likely.

## Introduction

Many studies have focused on the prevalence of dental pathology and wear due to the insight these variables can give into the health and lifestyle of past populations (e.g., Eshed et al., 2006; Hillson, 2001; Keenleyside, 2008; Krogman, 1940). When pathological frequencies are investigated on a population level it is possible to infer certain dietary and behavioral habits, particularly when evidence from isotopic, environmental, and cultural sources are also considered (Bonsall, 2014; Bonsall & Pickard, 2015; Lillie, 1996; Lillie & Richards, 2000; Littleton & Frohlich, 1993). To explore differences in dental pathologies at a population level for the present study, analyses were carried out on remains form a medieval cemetery from a port town in Southwest England. This location presents an opportunity to study diachronic changes a small geographical area. Previous studies provide comparative pathological and dietary information for Neolithic (Hedges et al., 2008), Roman (Cheung et al., 2012; Simmonds et al., 2008), and medieval populations (Dawson & Brown, 2011; Enright & Watts, 2002) from Gloucestershire and the town of Gloucester itself. Further comparisons can also be drawn with other medieval sites around the United Kingdom.

Literary and environmental sources suggest cereals were main components of the English medieval diet (Connell et al., 2012; Woolgar et al., 2006; Thomas et al., 1997). Wheat was used for bread making and oats in pottage, a basic broth consumed by most status groups. Cereals were also malted to produce ale and vineyards would have enabled the production of wine at some priories (Connell et al., 2012). Excavations at St Mary Spital, an urban site in London, uncovered the remains of fruits, nuts, vegetables, herbs and spices, providing evidence for a wide variety of foods within urban environments (Connell et al., 2012). Pulses and dairy products were also likely important dietary components (Wadsworth, 1992; Woolgar et al., 2006; Novak, 2015). Pork and lamb were often eaten but documentary sources suggest beef was most commonly consumed (Albarella, 2006). Marine fish was more commonly consumed due to restrictions on freshwater fishing; however, some higher status groups had private fishponds, which provided fresh water varieties to the select few (Schofield & Vince, 2003). The docks and location of the town of Gloucester would have enabled access to marine fish, as well as frequent trade at the marketplace, which led to a variety of foodstuffs being available to the population. The town was a strategically placed defense post along the River Severn and one of the wealthiest cities in England by medieval times.

The presence of dental caries is virtually universal among human populations, with frequency and location on the dentition varying by diet. Caries form when bacteria, such as *Streptococcus mutans* and *Lactobacillus acidophilus*, demineralize dental tissue through the release of acids when sugars and starches are metabolized (Larsen et al., 1991; Byun et al., 2004). These acids form active lesions as the oral pH level is lowered for extensive periods (Gussy et al., 2006). Some foods are more cariogenic than others, with those that contain high levels of refined carbohydrates and sugars particularly virulent. Foods that are linked with low rates of caries include tough and fibrous items, with a tendency to create a more alkaline oral environment due to high levels of saliva production (Prowse et al., 2008; Rohnbogner & Lewis, 2016; Moynihan, 2000). Meat and dairy products are also among the foods that have been associated with a low caries frequency (Novak, 2015). Past research into caries has similarly highlighted differences in prevalence when considering sex, age and social status (Larsen et. al., 1991; Walker, 1986; Rohnbogner & Lewis, 2016).

Dental calculus is a biofilm that forms when bacteria along with organic and inorganic material, adhere to a tooth surface as a plaque deposit that mineralizes over time (Lieverse, 1999). A variety of dietary factors can influence calculus formation on a population level, making its interpretation more complex than that for caries (Novak, 2015; Delgado-Darias et al., 2006; Lieverse, 1999). Further complications arise due to high rates of postmortem damage (Hillson, 2008). Diets with high levels of carbohydrates and proteins have been shown to have elevated frequencies of calculus (Littleton & Frohlich, 1989; Lillie, 1996; Lieverse, 1999; Lillie & Richards, 2000), but a diet high in carbohydrates and low in protein has been associated with high rates of both calculus and caries (Keenleyside, 2008; Šlaus et al., 2011).

Ante-mortem dental chipping and non-masticatory wear can give insight into cultural practices (Belcastro et al., 2007; Constantino et al., 2010; Scott & Winn, 2011). Cultural deviation causes differences in the position of chipping on dentitions. For example, tooth chipping in agriculturalists tends to be located on the anterior dentition, with hunter-gatherers more likely to present fractures on posterior teeth (Scott & Winn, 2011; Towle et al., 2017). Non-masticatory behavior has remained the focus of studies looking at chipping in more recent populations, due to the wide variety of chipping patterns represented (e.g., Bonfiglioli et al., 2004; Larsen, 2015; Lous, 1970; Molnar at al., 1972).

In this exploratory study the frequencies of different pathologies and types of wear present on the dentitions of a medieval cemetery population from Gloucester are investigated. In particular, through the examination of the rates of caries, calculus, antemortem chipping and non-masticatory wear, a greater understanding of the lifestyles and diets of this port town population can be provided. Given data is available in the literature on contemporary sites in other areas of the UK, as well as in the same area but for different time periods, this study aims to explore if this population retains a similar diet and behavior’s over time or if the time period involved is the principal predictor of pathology frequencies.

## Materials and methods

Dental pathologies were recorded in 41 adult individuals (914 permanent teeth) from the medieval cemetery of St Owens Church, in Southgate Street, Gloucester (Figure 1). Occupation of the area originally formed as a Roman suburb before becoming part of Llanthony Priory in 1137 (Rawes, 1990). The church was founded by the first sheriff of Gloucester, Roger de Pitres, who held the position between 1071 and 1082 (Herbert, 1988; Morris, 1918). The church was built outside the south gate of the city and served the parish of St Owens until 1643 when the area was cleared and fired prior to attack by the Royalist army during the siege of Gloucester. The parish of St Owens was merged with that of St Mary de Crypt in 1646. Although the Restoration in 1660 listed St Owens as a separate parish again, the church was never rebuilt, with the parishioners continuing to attend services at St Mary de Crypt (Herbert, 1988).

**Figure 1.**
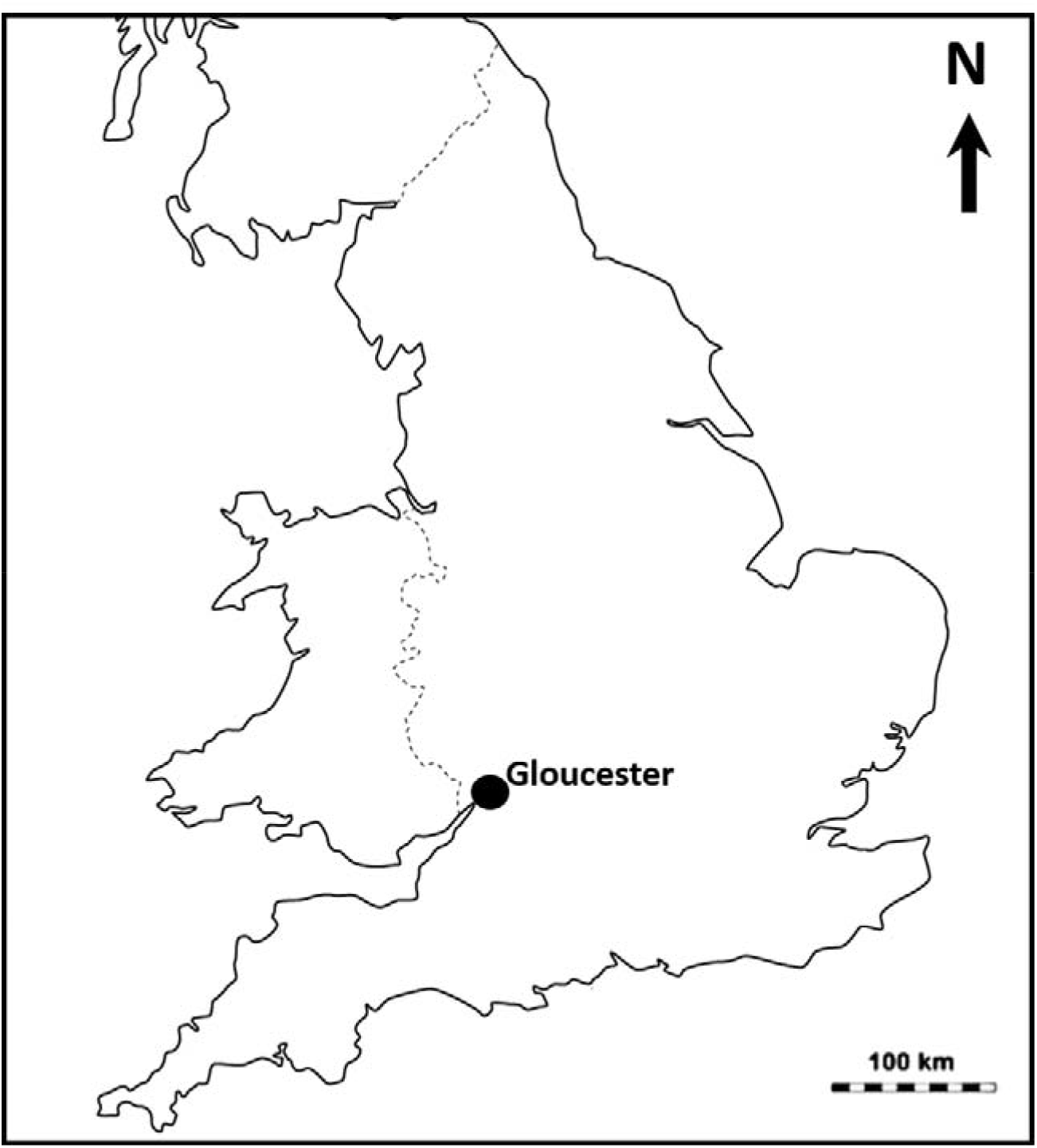
Map showing the location of Gloucester, UK.

A biological profile was carried out on each individual using a multivariate approach (Lovejoy et al., 1985). Age was assessed using up to six morphological techniques from the skeleton where the element was present (Brooks & Suchey, 1990; Işcan et al., 1984, 1985; Webb & Suchey, 1985; Meindl & Lovejoy, 1985). Sex was assessed using morphological techniques for the skull and pelvis (Bass, 1995; Phenice, 1969), along with metric measurements from the post-cranial remains (Stewart, 1979; Berrizbeitia, 1989; Dwight, 1984).

Each tooth was examined macroscopically under good lighting with a 10x hand lens used for clarifying certain pathologies, particularly caries and chipping. The following variables were focused on: calculus, caries, antemortem tooth loss, abscesses, and antemortem chipping. For each, the frequency was calculated as follows:

### [Total number of teeth displaying the marker/total number of teeth observed]*100

A carious lesion was recorded if there was clear cavitation, with color changes alone not recorded. The severity and position of lesions on a tooth were recorded. Caries severity was scored on a scale of 1 to 4 following Connell & Rauxloh (2003), with (1) enamel destruction only; (2) involvement of dentine but pulp chamber not exposed; (3) destruction of dentine with the pulp chamber exposed; (4) gross destruction with the crown largely destroyed. Lesion location was also recorded (e.g., distal, buccal, occlusal, lingual, mesial and gross). Calculus presence and severity were recorded according to the criteria of Brothwell (1981). The three categories recorded were (1) slight: minimal and straight line; (2) moderate: up to 50% of the tooth surface covered; (3) severe: more than 50% of the tooth surface covered. A tooth was recorded as lost antemortem if the alveolar socket contained signs of bone resorption (Ortner, 2003; Novak, 2015). Therefore, if a tooth was not present but the alveolar socket contained no sign of remodelling the tooth was recorded as lost post-mortem. Alveolar abscesses have been often confused with other voids in the mandible and maxilla (Ortner, 2003; Dias & Tayles, 1997; Ogden, 2008; Hillson, 2005). In this study, an abscess was recorded if a smooth-walled sinus exited from the cortical bone of an associated tooth. Dental attrition was marked according to Smith (1984) for anterior teeth on a scale of 1 to 8 and Scott (1979) for molars, on a scale of 1 to 10. Notches and grooves on the dentition, caused by non-masticatory cultural activity, were also recorded.

The interaction between different dental pathologies is often complex. It has been suggested that because caries can lead to antemortem tooth loss, a sample may not give a true representation of the overall effect of caries on that population. This possibility led to a number of proposed methods that help counteract any association (e.g., Kelley et al., 1991; Lukacs, 1995; Duyar & Erdal, 2003). However, this study followed Meinl et al. (2009) by not using corrective methods, and instead displaying AMTL frequencies separately as an independent factor. This approach allowed direct comparisons with other population samples.

## Results

Table 1 presents per tooth frequencies for different pathologies divided by tooth type, jaw, sex and age. Abscessing was found in 2.55% of alveolar bone associated with a tooth, with 14 individuals having at least one abscess (34.15% of individuals). The prevalence of ATML is 6% for the sample as a whole. Dental wear was unexceptional, with older individuals commonly displaying much occlusal enamel. The mean wear score for anterior teeth, here including premolars, is 3.5 and molars 4.4 (Table 2). 17.6% of posterior teeth quadrants, and 12% of anterior teeth have a wear score of 6 or above. Overall, mandible molars were more worn than maxilla, with buccal quadrants of mandibular teeth having higher wear than lingual (means: 4.80 vs. 4.48), whilst the opposite is true for maxillary teeth (means: 3.91 vs. 4.36 respectively).

**Table 1.** Per tooth percentage (%) frequencies for each pathological condition present on the dentition; split by sex, age and tooth type.

| Pathology | Sample grouped by |  |  |  |  |  |  |  |  |
| --- | --- | --- | --- | --- | --- | --- | --- | --- | --- |
|  | All Teeth | Position of dentition |  | Skeletal location |  | Sex |  | Age |  |
|  |  | Anterior | Posterior | Maxilla | Mandible | Male | Female | < 35 yrs | > 35 yrs |
| Caries | 23.97 | 10.96 | 32.21* | 23.96 | 23.97 | 25.33 | 28.71 | 20.63 | 32.93* |
| Calculus | 74.3 | 79.66 | 70.89* | 70.74 | 77.50* | 82.37 | 77.45 | 71.11 | 82.93* |
| Chipping | 6.7 | 10.14 | 4.49* | 7.03 | 6.39 | 9.38 | 4.29* | 4.93 | 11.52* |
| Abscess | 2.55 | 1.87 | 3.01 | 3.45 | 1.78 | 1.97 | 4.38 | 1.46 | 5.51* |
| AMTL | 6.06 | 2.13 | 8.67* | 5.75 | 6.34 | 5.47 | 9.69* | 3.94 | 11.81* |

\*Chi-square significant difference between variables at 0.05 level

**Table 2.**
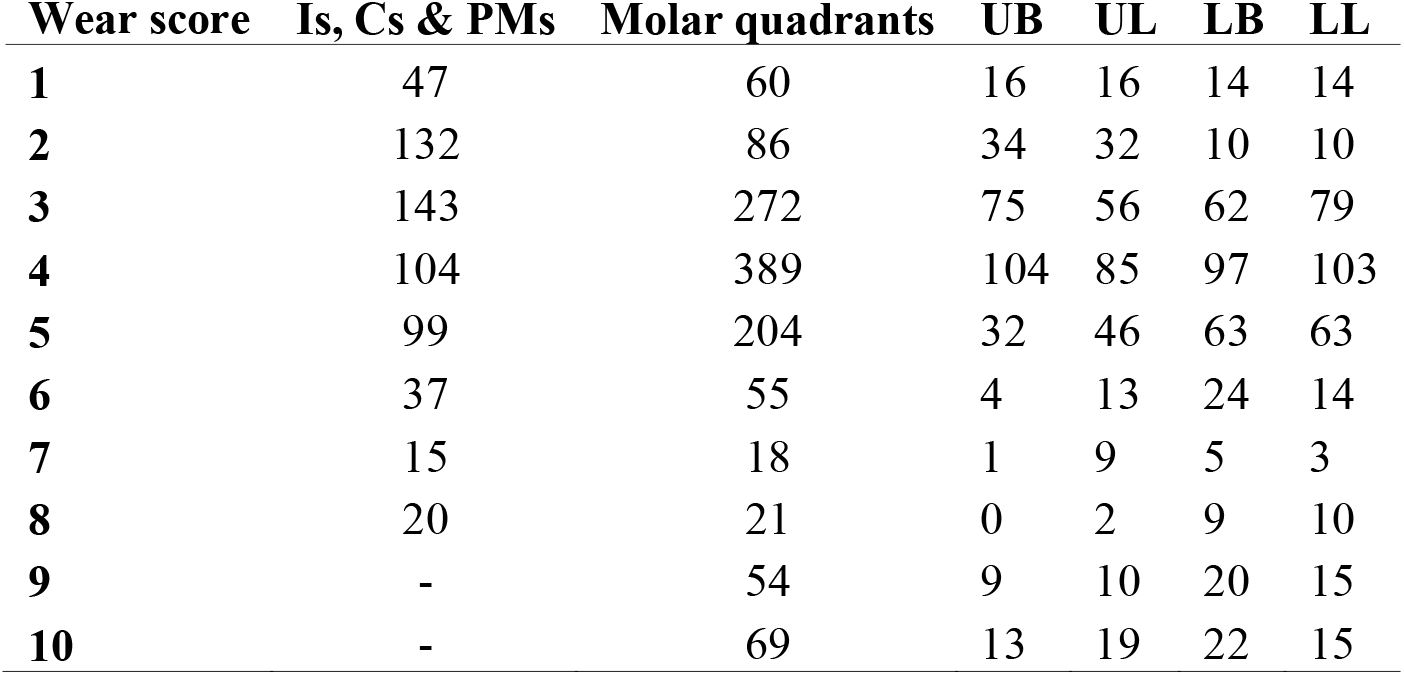
Wear scores for individual teeth. Molar wear is based on Scott (1979), with teeth split into quadrants. The two buccal and lingual quadrants are combined to give UB (upper buccal), UL (upper lingual), LB (lower buccal), and LL (lower lingual). All other teeth are recorded following Smith (1984), with Is, Cs & PMs: incisors, canines and premolars respectively.

Nearly 88% of individuals in the sample exhibit at least one carious tooth, with 23.97% of all teeth affected (n=220/918). Lesions are most common on interproximal surfaces (mesial: 26%; distal: 25%) followed by occlusal (19%) and buccal/labial (15%) surfaces (Figure 2). Figure 3 highlights the differences in severity of caries and calculus between sex and age. Both sexes had similar proportions of caries in all four severity categories, with females exhibiting a slightly higher frequency of gross caries; however, this difference is not statistically significant (χ^2^= 0.542, 1 df, p= 0.4616). There are 259 carious lesions across the sample, with 39 teeth having two or more lesions (4.25%). Only five individuals were caries-free, meaning that 87.8% of individuals had at least one carious lesion. Posterior teeth were significantly more affected than anterior (χ^2^= 54.016, 1 df, p>0.05), with 32.21% and 10.96% of teeth affected respectively. Calculus was also common throughout the sample with 74% of teeth affected; of these, 5.4% were severely affected (Brothwell grade 3; Table 1). Only one individual did not evidence calculus, although this individual had multiple antemortem teeth lost and gross caries.

**Figure 2.**
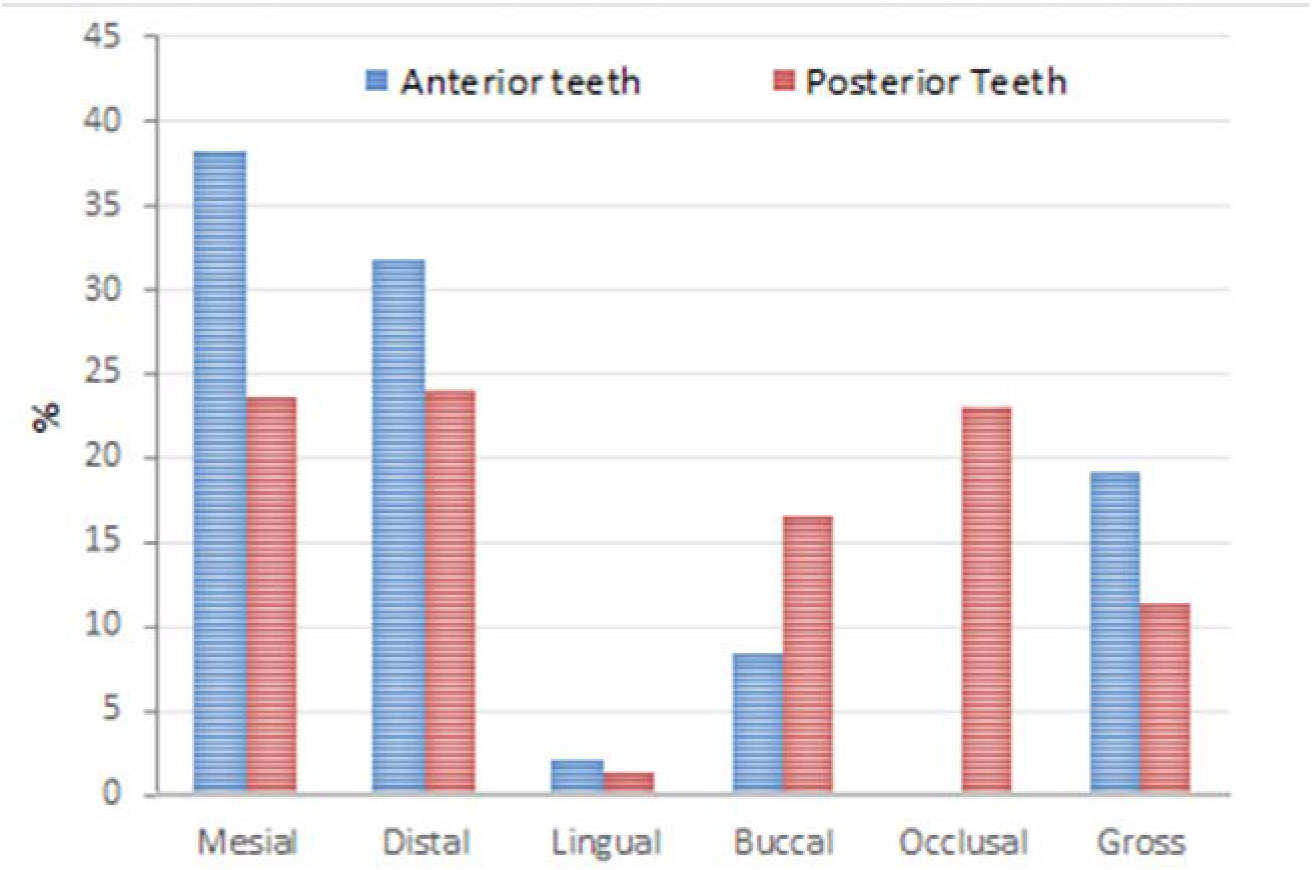
Position of carious lesions as a percentage of teeth with caries. Teeth are split into anterior and posterior.

**Figure 3.**
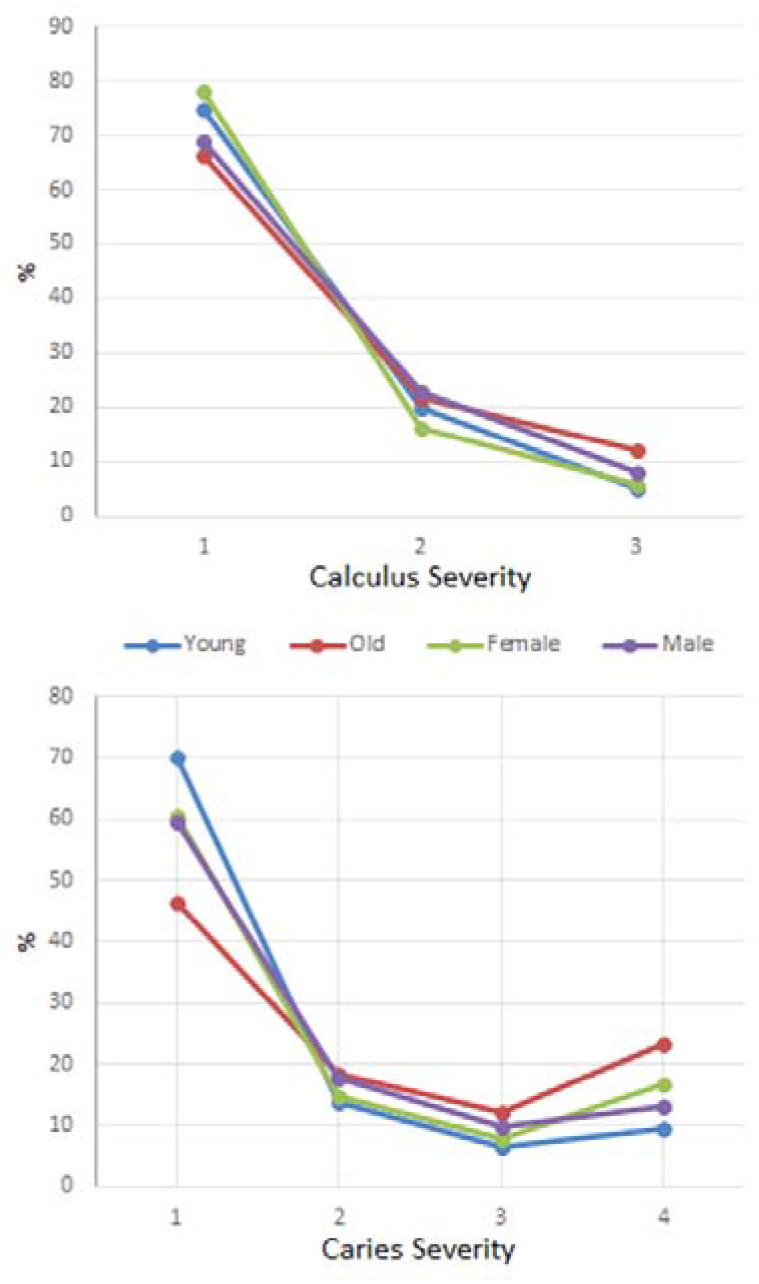
Per tooth caries and calculus severity split by age and sex with each severity grade being a percentage of all teeth with the pathology. Caries severity is based on criteria following Connell & Rauxloh (2003), and calculus Brothwell (1981).

All pathologies more commonly affected older individuals (Table 1). Overall caries, calculus, and abscess rates are not significantly different between the sexes; however this was not the case for chipping and ATML, as males present higher rates of chipped teeth (χ^2^= 6.885, 1 df, p>0.05), and females more ATML (χ^2^= 5.005, 1 df, p>0.05). A significant difference was also found between maxilla and mandible for the presence of calculus, where 77.50% of mandibular teeth and 70.74% of maxillary teeth were affected. Calculus and chipping are more common on anterior teeth, whereas caries and ATML are more evident on posterior teeth (Table 1).

Non-masticatory cultural notches are visible on eight maxillary central incisors from six individuals, i.e., two males, two females, and two for which sex could not be assessed. These notches are small, all graded as a 1 or 2 on the Bonfiglioli (2004) three-point severity scale.

## Discussion

The frequency of antemortem tooth loss in the St Owens population sample is similar to other medieval collections around Europe (e.g., Novak, 2015; Slaus et al., 1997; Vodanović et al., 2005; Caglar et al., 2007). The rate of AMTL is not particularly high for this period, considering the high rates of caries, which is likely due to other factors, including periodontitis, fractures, and gross attrition (Caglar et al., 2007; Esclassan et al., 2009; Whittaker et al., 1998). However, the significantly higher rate of ATML in females is most likely due to caries, which is supported by the increased abscess and gross caries frequencies relative to males. Previous research also found higher rates of caries in females, likely related to dietary and behavioral differences (Lukacs, 1996; Walter et al., 2016).

Non-masticatory dental wear is relatively common in archaeological samples (e.g., Irish & Turner, 1997; Turner et al., 2003; Bonfiglioli et al., 2004; Scott & Winn, 2011). Notches in this sample are in individuals of all age and sex group, which relate to occupation or habitual behaviours. Gloucester was a port town, with a thriving economy. The range of occupations that may have contributed to habitual behaviours causing this tooth wear include haberdashers, weavers, shoemakers and fishermen. Aside from merchants dealing in the import and export of goods, there would have been workforce preparing the goods for sale. Similarly, higher rates of chipping on anterior teeth is suggestive that many of these fractures were the result of food processing or, more likely, other cultural/social behavior (Belcastro et al., 2007; Constantino et al., 2010; Bonfiglioli et al., 2004). The ratio of anterior to posterior chips is similar to that in other European samples from this time, in which food preparation and tool use have been proposed as likely causes (Scott & Winn, 2011; Belcastro et al., 2007). That said, the overall chipping rate is still low compared to some samples in which teeth were clearly used more often for non-masticatory activities (e.g., Scott & Winn, 2011; Turner & Cadien, 1969). Males have significantly more chips per dentition than females, which may relate to using their teeth more for non-masticatory activities.

Generally, it is thought that most medieval populations had high rates of wear, particularly compared to modern samples (Esclassan et al., 2009; Hillson, 1996; Srejić, 2001). It was suggested by d’Incau and Rouas (2003) that difference could be due to consuming large amounts of abrasive food and intense masticatory pressures required during eating. Hard particles incorporated into the diet direct from foods, or from food processing, have also been suggested (Esclassan et al., 2009). The lack of severe wear in most individuals at this site and presence of moderate wear, in even the oldest individuals, suggests their diet may not have been as abrasive at other contemporary sites. Additionally, the few antemortem dental chips on posterior teeth suggests hard objects, especially larger ones, were not regularly consumed (Scott & Winn, 2011; Bonfiglioli et al., 2004).

The caries rate is far higher than in other European medieval population samples. It has often been noted that the caries rate in medieval Europe varied widely, with assemblages ranging from 3% to 17.5% of teeth affected (Novak, 2015; Esclassan et al., 2009; Vodanović et al., 2005; Caglar et al., 2007; Meinl et al., 2010; Pap, 1986; Manzi et al., 1999; Belcastro et al., 2007; Slaus et al., 2010). Luis (2007) found that rates in late medieval South East England vary between 4% and 15%. Roman samples are similarly affected, i.e., 5.9% to 10.6%. These values fit with the interpretation that caries was prevalent in Roman Britain, falling to a low by the early medieval and increasing again in late medieval times (Roberts & Cox, 2003). Caries frequency for this Gloucester sample is more like those of later sites (i.e. post-medieval) in Great Britain (Mant & Roberts, 2015). An increase in consumption of highly cariogenic foods, perhaps facilitated through being a port town, likely explains this high rate of lesions.

Although the frequency of caries is high in this sample, the location of lesions is similar to contemporary populations, with posterior teeth and interproximal areas most affected (Slaus et al., 1997, 2011; Vodanović et al., 2005; 2012; Watt et al., 1997; Manzi et al., 1997; Varrela, 1991; Esclassan et al., 2009; Srejić, 2001). Groups with higher attrition rates caused by abrasive and tough foods tend to have few occlusal caries and a far higher proportion of interproximal lesions (Maat & Van der Velde, 1987; Meinl et al., 2009). In contrast, populations with a high intake of refined carbohydrates and sugars tend to show high frequencies of occlusal caries (Hillson, 1996). This variation fits with the fact that although interproximal areas are most affected, caries on occlusal surfaces are higher than at other medieval sites (Novak, 2015). The fact that interproximal areas are still most affected is likely explained by higher attrition than in later times, but may be influenced by the high frequency of calculus-facilitated caries formation in these areas (Tomczyk et al., 2013).

The presence of calculus is harder to interpret in terms of diet (Delgado-Darias et al., 2006). Although oral hygiene, salivary flow and other non-dietary factors can influence calculus rates (Lieverse, 1999), diet is the main factor affecting frequencies on a population level (Novak, 2015). This relationship is far from clear, however, with diets both high in carbohydrates as well as proteins showing similarly high levels of calculus (Littleton & Frohlich, 1989; Lillie, 1996; Littleton & Frohlich, 1989; Lieverse, 1999). Calculus frequency varies drastically among medieval sites in Europe, from 27% to almost 90% (Manzi et al., 1999; Belcastro et al., 2007; Slaus et al., 2010; Delgado-Darias et al., 2006; Vodanovic et al., 2012). Novak (2015) notes a very high rate in a large sample of medieval Irish from different sites, but unlike this study the rate of caries is low. Novak (2015) suggests both can be explained by incorporation of carbohydrates and dairy products. Therefore, frequent caries in the Gloucester sample suggest dairy products were likely not relied upon heavily. This occurrence is greater than at most other British sites of this age yet, interestingly, it is similar to values for earlier and contemporary Gloucester (Roman Gloucester: 66.7% of teeth; 3.2% at grade 3; Simmonds et al., 2008). This agrees with research that suggests during the Roman period different provinces differed markedly in diet, mostly notably with Southwest populations likely opposing new plant foods (van der Veen et al., 2008; Rohnbogner & Lewis, 2016). This supports the conclusion that populations in and around Gloucester were likely also consuming a diet substantially different from other areas of the UK during the medieval period and this may reflect difference in foods available but also potentially local cultural and cookery practices.

In sum, a diet high in particularly cariogenic carbohydrates is likely the cause for high rates of calculus and caries in this sample (Keenleyside, 2008; Šlaus et al., 2011). Because these rates are also high in the area during the Roman period may suggest a continuation of certain foods or behavioral practices or, more likely, Gloucester was a key port town throughout. The conclusion that Gloucester was a prosperous town with a diet high in ‘luxury’ foods such as meat and sugary carbohydrates is also supported by recent isotope analysis for this material (Chamberlain et al., 2016; Poulson et al., 2016).* Results from the present research and other multi-site analyses reveal that individual sites can vary dramatically during a given time particularly, as seen here, in the medieval period.

*Footnote. As part of a separate study, dental calculus samples from St Owens (n=26) were sent for carbon and nitrogen stable isotope testing at the University of Nevada, Reno (Chamberlain et al., 2016; Scott and Poulson, 2012; Poulson et al., 2013; 2016). The mean δ^13^C values (−21.34 ±0.79‰) for the St Owens chapel calculus samples were consistent with a diet focused on C3 plants, which was supported by documented evidence of trade and agriculture in Gloucester during the medieval period (Herbert, 1988). Mean δ^15^N values (12.53 ± 0.19‰) are consistent with a 1-2 level trophic shift (6.72‰) above the herbivore baseline (5.81 ± 0.19‰, n=51), indicating that the diet contained animal and marine sources, along with regular intake of omnivore or freshwater fish protein.

## Acknowledgments

We would like to thank the Museum of Gloucester for access and support in studying the skeletal collection housed at Liverpool John Moores University. The authors would also like to thank Richard Scott and Simon Poulson for making available the isotope data for use as a comparison. This research was supported by a studentship to the first author from Liverpool John Moores University.

